# The foot fault scoring system to assess walking adaptability in rats and mice: a reliability study

**DOI:** 10.1101/2020.11.03.366351

**Authors:** Lucas Athaydes Martins, Aniuska Schiavo, Léder Leal Xavier, Régis Gemerasca Mestriner

## Abstract

The foot fault scoring system of the ladder rung walking test is used to assess walking adaptability in rodents. However, the reliability of the ladder rung walking test foot fault score has not been properly investigated. This study was designed to address this issue.Two independent and blinded raters analyzed 20 rat and 20 mice videos. Each video was analyzed twice by the same rater (80 analyses per rater). The intraclass correlation coefficient (ICC) and the Kappa coefficient were employed to check the accuracy of agreement and reliability in the intra- and inter-rater analyses of the ladder rung walking test outcomes. Excellent intra- and inter-rater agreement was found for the forelimb, hindlimb and both limbs combined in rats and mice. The agreement level was also excellent for total crossing time, total time stopped and number of stops during the walking path. Rating individual scores in the foot fault score system (0 to 6) ranged from satisfactory to excellent, in terms of the intraclass correlation indexes. Moreover, we showed experienced and inexperienced raters can obtain reliable results if supervised training is provided. We conclude the ladder rung walking test is a reliable and useful tool to study walking adaptability in rodents and can help researchers address walking-related neurobiological questions.

## 1. INTRODUCTION

Walking adaptability can be defined as a complex sensory-motor function, qualified or required to control and coordinate various degrees of freedom in joints, in a variety of environmental contexts, or that interfere with locomotion [1-3]. Gait is influenced by the temporal and spatial integration of the cognitive and neuromusculoskeletal neural systems [4]. Moreover, the ability to adapt gait according to environmental context is a crucial aspect in maintaining body stability and preventing falls [5-8].

Whilst several studies into walking adaptability have focused on human biomechanics [2, 9, 10], animal models can usefully provide neurobiological insights at the cellular and molecular level [11-13]. For instance, the Ladder Rung Walking Test (LRWT) has been used to assess walking adaptability [14, 15] in unilateral ischemic injury in the motor cortex [12, 16]; spinal cord injury [17, 18]; dopaminergic depletion induced by 6-hydroxydopamine (a model of Parkinson’s disease) [19]; neonatal white matter injury [20] and stress-related conditions [7, 13, 21].

The LRWT can assess walking patterns by using measures of inter-foot coordination, foot support, fore and hindlimb kinematics, step and gait cycles, gait speed, and the ability to adapt walking by applying a foot-fault score [16, 17]. The test provides measures of gait adaptability with emphasis in forelimb and hindlimb function by applying the foot-fault score [15]. The foot-fault score system is widely used in the literature since it requires only a hand camera and a minimally trained researcher to analyze the video and apply the foot-fault score [13, 14]. This method may avoid common pitfalls that occur when using reflective markers on the flexible skin of rodents [22, 23] and gives a measure of the success in adapting walking [7, 13].

The foot-fault score system is a 7-point category scale in which the quality and appropriateness of foot placement is judged by analyzing a video recording, frame-by-frame, of rodents walking along a 1-meter long horizontal ladder. The rungs are arranged in a pattern that requires murine ability to adapt walking [14, 15]. However, to the best of our knowledge, this test has not been properly assessed regarding its intra-rater and inter-rater reliability and reproducibility, which is a source of uncertainty. Current studies usually elect a single rater to analyze all videos in an attempt to minimize bias, which is scientifically insufficient. The present study sought to provide scientific information regarding the external validity of the LRWT findings in rodents, thus contributing to advancements in the field of neurobiology of walking adaptability.

## 2. MATERIALS AND METHODS

We randomly select 40 video recordings of rodents from our lab database (20 recordings of Wistar CrlCembe:WI rats and 20 of C57BL/6JUnib mice), that performed the horizontal ladder rung walking test. At the time of the original experiments, the animals were provided by the Center for Experimental Biological Models (CeMBE) of the Pontifical Catholic University of Rio Grande do Sul. The animals were housed in cages each containing three to four rodents on a 12-hour dark-light cycle with food and water available *ad libitum*, at a temperature of 22 to 24 °C. The experiments were carried out in accordance with the National Council for Animal Control and Experimentation (Concea) and all the procedures were approved by the University Animal Ethics Commission (CEUA) under protocol numbers 15/00442 and 15/00475.

### 2.1 Ladder rung walking test

We used two LRWT apparatus, one for rats and another adapted for mice. Both consisted clear Plexiglas side walls (100 cm long and 20 cm high). The diameter of the metal rungs varied, being 3 mm for rats and 2 mm for mice. The minimum and maximum gaps between the rungs also varied, being from 1 to 5 cm for rats and from 0.5 to 2.5 cm for mice. In both cases, the ladders were elevated horizontally 30 cm above the ground, with a neutral cage placed in the starting position and the animal’s home cage placed at the opposite end of the ladder (Figure 1). The between-wall distance was adjusted leaving 1 cm wider than the size of rodent to prevent the animal turning around during the crossing [13, 14, 24].

**Figure 1.**
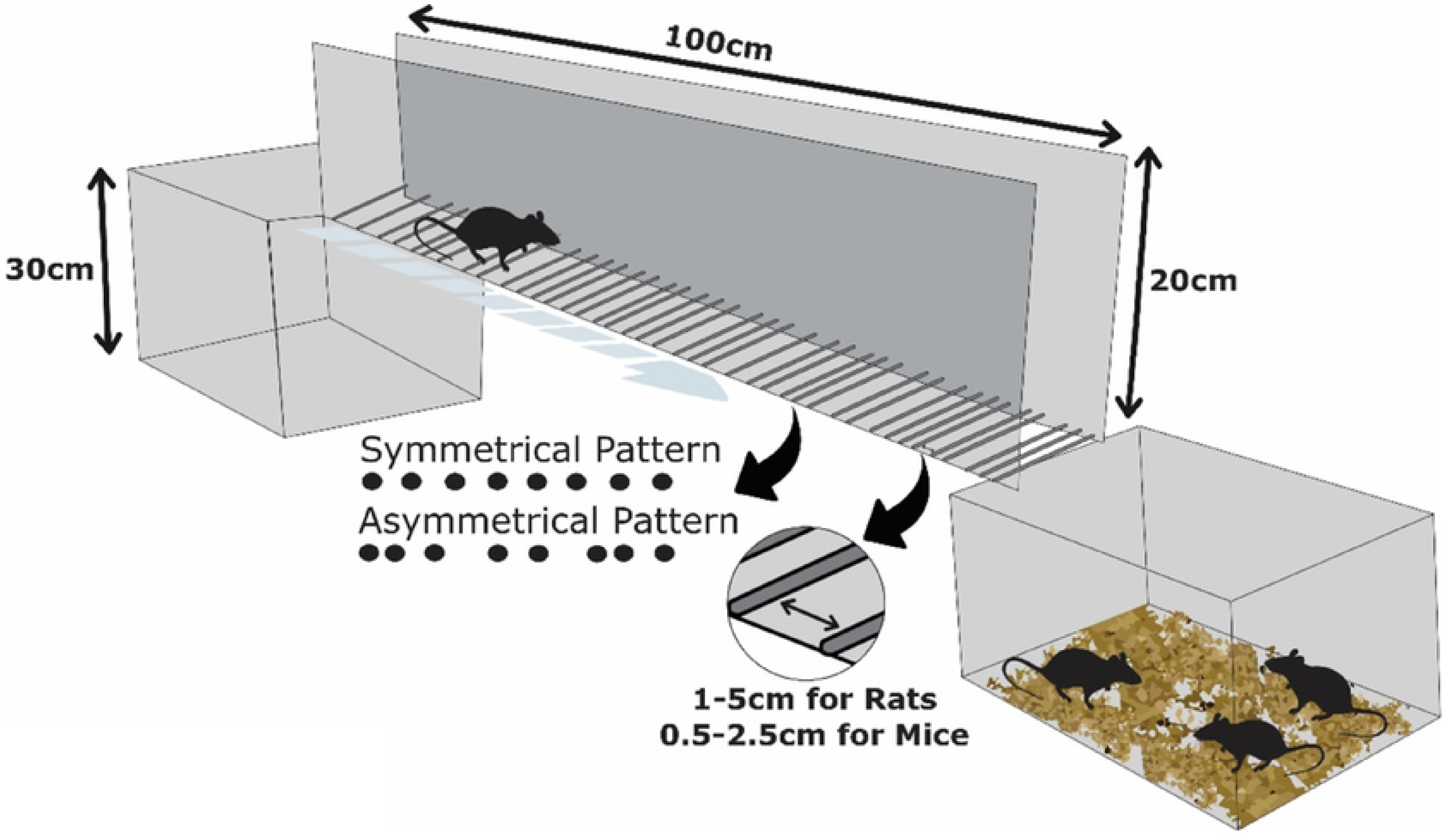
Schematic illustration of the ladder rung walking test apparatus.

The pattern of the metal rungs demands different degrees of walking adaptability and can be used to vary the complexity of the test. A regular arrangement allows animals to learn the position of the rungs over training sessions and to anticipate limb placement (Figure. 1, Symmetrical Pattern). In irregular patterns, rungs are randomly repositioned in each trial to prevent the rodents learning the rung sequence. Thus, irregular patterns are more useful when studying walking adaptability (Figure. 1, Asymmetrical Pattern) [7, 13, 14]. In this study, only irregular rung patterns were analyzed.

In the test, the animals were placed at the beginning of the ladder, walked along it, adapting their foot placement on the rungs until reaching the home cage (Figure 1). While performing the test, we filmed the rodents using a camera (GoPro Hero 4, 12 megapixels). An acquisition rate of 240 frames per second (FPS) in a lateral view was adopted allowing a *post-hoc* frame-by-frame video analysis.

### 2.2 Foot Fault Scoring System

To assess the fore and hindlimb placement on the rungs, which requires precise and coordinated foot positioning as well as stride and inter-limbic coordination a quantitative foot fault scoring system [14] derived from a categorical analysis was used. In the system, a frame-by-frame video recording analysis is performed to identify the steps in each limb and qualify foot placement using a 7-point category scale [14, 15] (Table 1). The score 0 is given when the limb did not touch the rung (missed a rung) and resulted in a fall (total miss). A fall is considered when the limbs fell between rungs and the animal’s posture and balance are disturbed. Score 1 is given when the limb slipped off a rung and a fall occurred (deep slip). Score 2 is given when the limb slipped off a rung during weight bearing, but a fall did not occur and the rodent interrupts walking (slight slip). Score 3 is given when, before weight bearing the limb on a rung, the rodent quickly lifted and placed the foot on another rung (replacement). Score 4 occurs when the limb is clearly about to be placed on a rung, but the rodent quickly changes the feet placement to another rung without touching the first rung (correction). Score 4 is also given when the limb is placed on a rung, but the animal removes the foot and repositions it on the same rung. Score 5 is given when the limb is placed on the rung either using the wrist or digits for the forelimb or heel or toes for the hindlimb (partial placement). Finally, score 6 is given when the full body weight bearing is applied on a rung with the midportion of the foot (correct placement) (Table 1).

**Table 1.** Rating scale for foot placement in the LRWT.

| Category | Type of foot misplacement |
| --- | --- |
| 0 | Total miss |
| 1 | Deep Slip |
| 2 | Slight Slip |
| 3 | Replacement |
| 4 | Correction |
| 5 | Partial Placement |
| 6 | Correct Placement |

The score given in each category is then multiplied by the frequency of foot placements in the same category. Afterwards, the sum of all the categories provides the total combined score (sum of the forelimb plus the hindlimb scores). The fully explained video protocol and all technical details to apply the foot fault score were previously published by Metz and Whishaw (2009).

In this study the following outcomes in the LRWT were assessed for inter-rater and intra-rater agreement: Total Crossing Time, Number of Stops, Total Time Stopped, Scores 0 to 6 for forelimb, Total Score for forelimb, Scores 0 to 6 for hindlimb, Total Score of hindlimb and the Combined Total Score of limbs.

The skilled walking performance score (SWPS) was represented as a percentage of the maximum possible performance (100%) *. The number of cycles (NC) each rodent took to cross the ladder was multiplied by 6 (the maximum score for each cycle in the foot fault score system) and the resulting number was considered the maximum possible performance of each animal in a trial (100%). Then, during a trial, each cycle was rated according to the foot fault score system and the sum of the obtained scores provided the total score in the trial (TS). Finally, the SWPS was represented as a percentage of the maximum possible performance (100%) [7, 25], as follows:

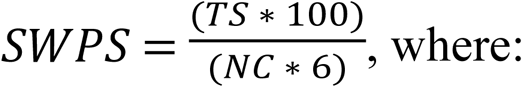

*SWPS* = skilled walking performance score

*TS* = total score in the trial

*NC*: number of cycles

6: maximum score for each cycle in the foot fault score system

### 2.3 Foot placement reliability between inter- and intra-rater

In order to asses inter- and intra-rater reliability, two independent and blinded raters (called I and II) analyzed 20 rat and 20 mice videos. Each video was analyzed twice by the same rater (80 analyses per rater). The videos were named randomly by another independent researcher (not involved in the analyses) to prevent raters I and II from perceiving half of the videos were the same. Thus, each video had a different number to ensure a blinded reproducibility analysis. Rater I (Schiavo, A) was inexperienced in the foot fault score and received supervised training before starting data collection. Rater II (Martins, LA) had previous experience and publications using the LRWT [7, 13].

### 2.4 Statistical Analysis

Descriptive statistics were used to characterize the sample profile in the SWPS. The intraclass correlation coefficient (ICC) and the Kappa coefficient were employed to verify the accuracy of agreement and reliability in the inter-rater and intra-rater analyses of the foot fault scores. Agreement values in ICC greater than 0.75 were considered “excellent”; between 0.4 and 0.75 “satisfactory” and those <0.4 were considered “poor”. When negative ICC values (difference between values greater than sample variance) occurred, the data were replaced by zero [26, 27]. The statistical analysis was performed using the software Statistical Package for the Social Sciences (SPSS) 20.0.

## 3. RESULTS

### 3.1 Inter-rater reliability for rat

The LRWT analyses in rat demonstrated rater I and II achieved an excellent agreement in the combined total score of limbs (ICC=0.938/p=0.0001). Regarding all the timed outcomes, the total crossing time (ICC=0.994/p=0.0001) and the total time stopped (ICC=0.992/p=0.0001) agreement levels were considered excellent, as were the variable number of stops (ICC=0.957/p=0.0001). Thus, the reliability between the total score for forelimb and hindlimb placement was shown to be excellent.

Furthermore, we analyzed the reliability among all scores described in the test, specifically, in the categories 0 to 6 for each of the limbs evaluated. For the forelimb, the data showed an excellent reliability in scores 0, 1 and 2, varying from ICC 0.839 to 1 (p=0.0001) as well as for scores 5 and 6 (ICC 0.813 and 0.854, respectively / p=0.0001). However, for the forelimb scores 3 and 4, the raters obtained a satisfactory agreement (ICC 0.721 and 0.551, respectively / p≤0.045). Similarly, for the hindlimb excellent reliability was obtained for scores 0, 1, 3 and 4, with the ICC ranging from 0.889 to 0.931 (p=0.0001). The reliability for scores 2, 5 and 6 was also considered satisfactory (Table 2).

**Table 2.** Agreement between raters I and II regarding the outcomes obtained in the LRWT in rat.

| **Foot fault scoring in rat** |  |  |  |
| --- | --- | --- | --- |
| **Outcome** | **ICC (IC95%)** | **Cronbach's Alpha** | **p Value** |
| Combined Total Score | 0.938(0.844-0.976) | 0.938 | 0.0001* |
| Total Crossing Time | 0.994(0.985-0.998) | 0.994 | 0.0001* |
| Number of Stops | 0.957(0.892-0.983) | 0.957 | 0.0001* |
| Total Time Stopped | 0.992(0.980-0.997) | 0.992 | 0.0001* |
| **Forelimb Placement** |  |  |  |
| **Outcome** |  |  |  |
| Score 0 | 1(1-1) | 1 | 0.0001* |
| Score 1 | 0.839(0.594-0.936) | 0.839 | 0.0001* |
| Score 2 | 0.903(0.754-0.961) | 0.903 | 0.0001* |
| Score 3 | 0.721(0.295-0.889) | 0.721 | 0.004* |
| Score 4 | 0.551(0.000-0.822) | 0.551 | 0.045* |
| Score 5 | 0.854(0.631-0.942) | 0.854 | 0.0001* |
| Score 6 | 0.813(0.528-0.926) | 0.813 | 0.0001* |
| Total Score | 0.879(0.695-0.952) | 0.879 | 0.0001* |
| **Hindlimb Placement** |  |  |  |
| **Outcome** |  |  |  |
| Score 0 | 0.889(0.719-0.956) | 0.889 | 0.0001* |
| Score 1 | 0.931(0.826-0.973) | 0.931 | 0.0001* |
| Score 2 | 0.593(0.000-0.839) | 0.593 | 0.028* |
| Score 3 | 0.889(0.719-0.956) | 0.889 | 0.0001* |
| Score 4 | 0.889(0.719-0.956) | 0.889 | 0.0001* |
| Score 5 | 0.41(0.000-0.620) | 0.41 | 0.456 |
| Score 6 | 0.592(0.000-0.839) | 0.592 | 0.029* |
| Total Score | 0.931(0.826-0.973) | 0.931 | 0.0001* |

\* Statistically significant difference.

The individual results for each animal in relation to SWPS are shown in Figure 2. In addition, the frequency of each score (1 to 6) for hindlimb and forelimb of each rodent is shown in Figure 3, 4 and 5.

**Figure 2.**
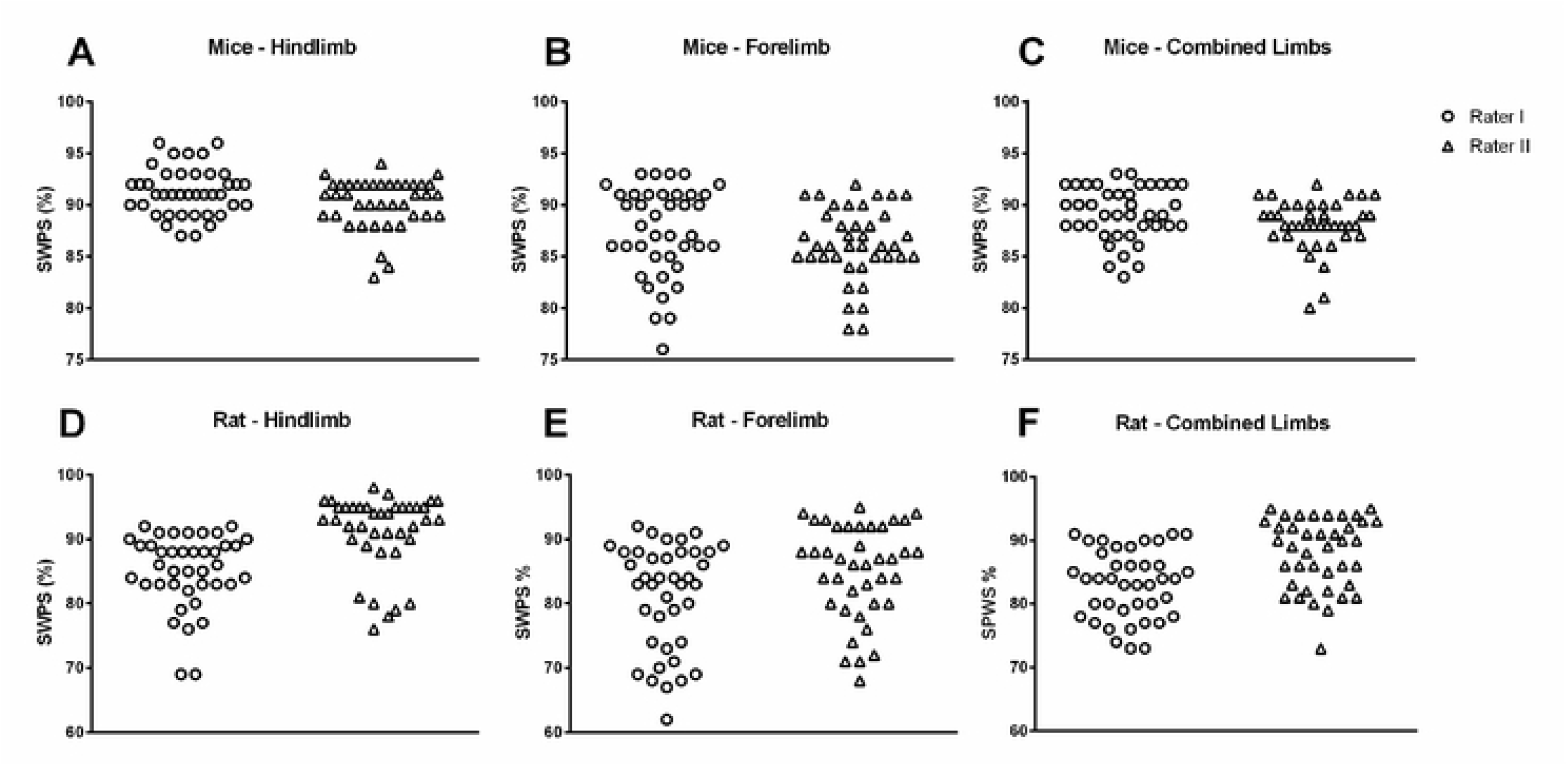
SWPS obtained by Rater I and Rater II.

**Figure 3.**
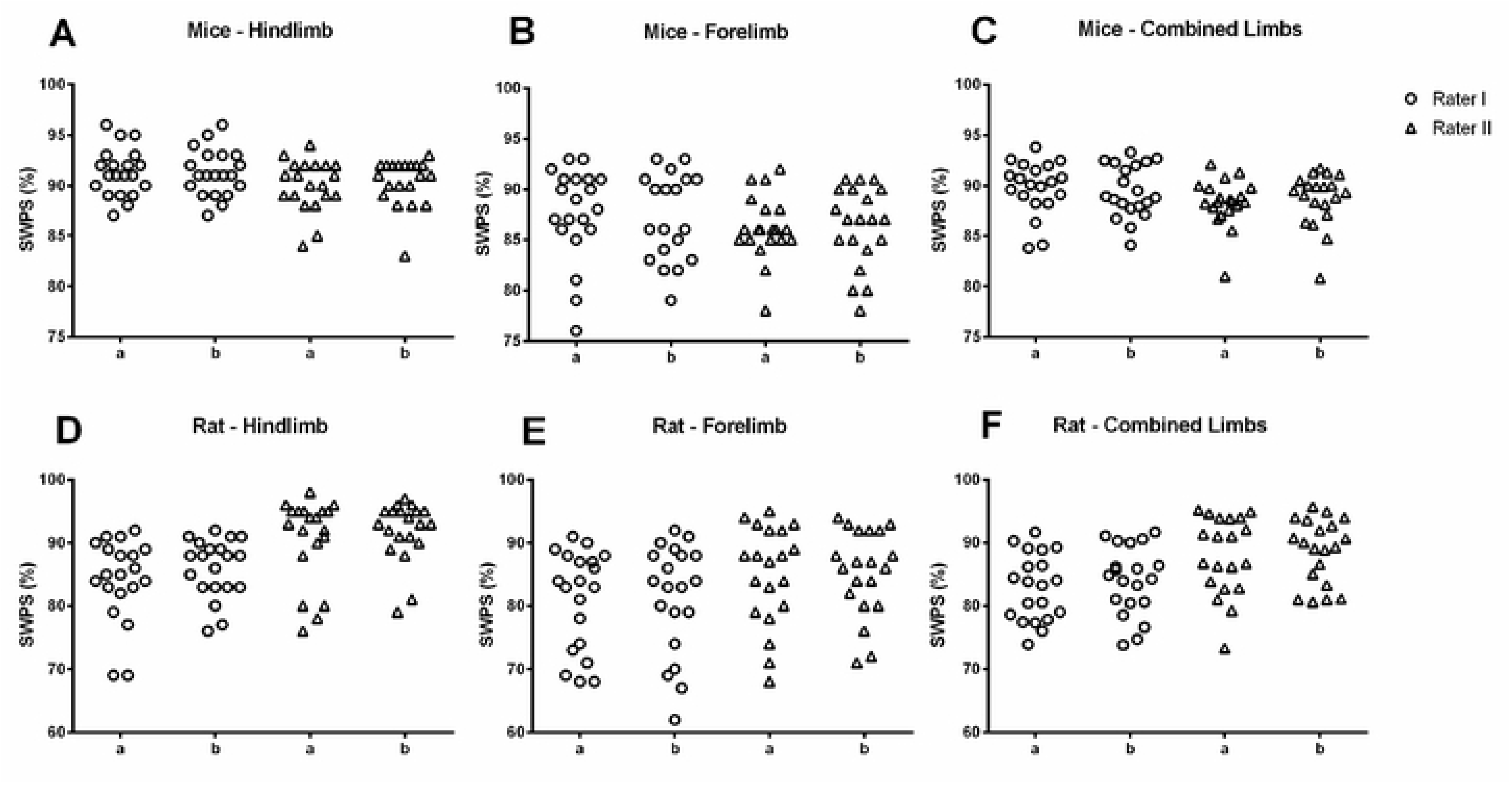
SWPS obtained at first (a) and second assessment (b) by Rater I and Rater II.

**Figure 4.**
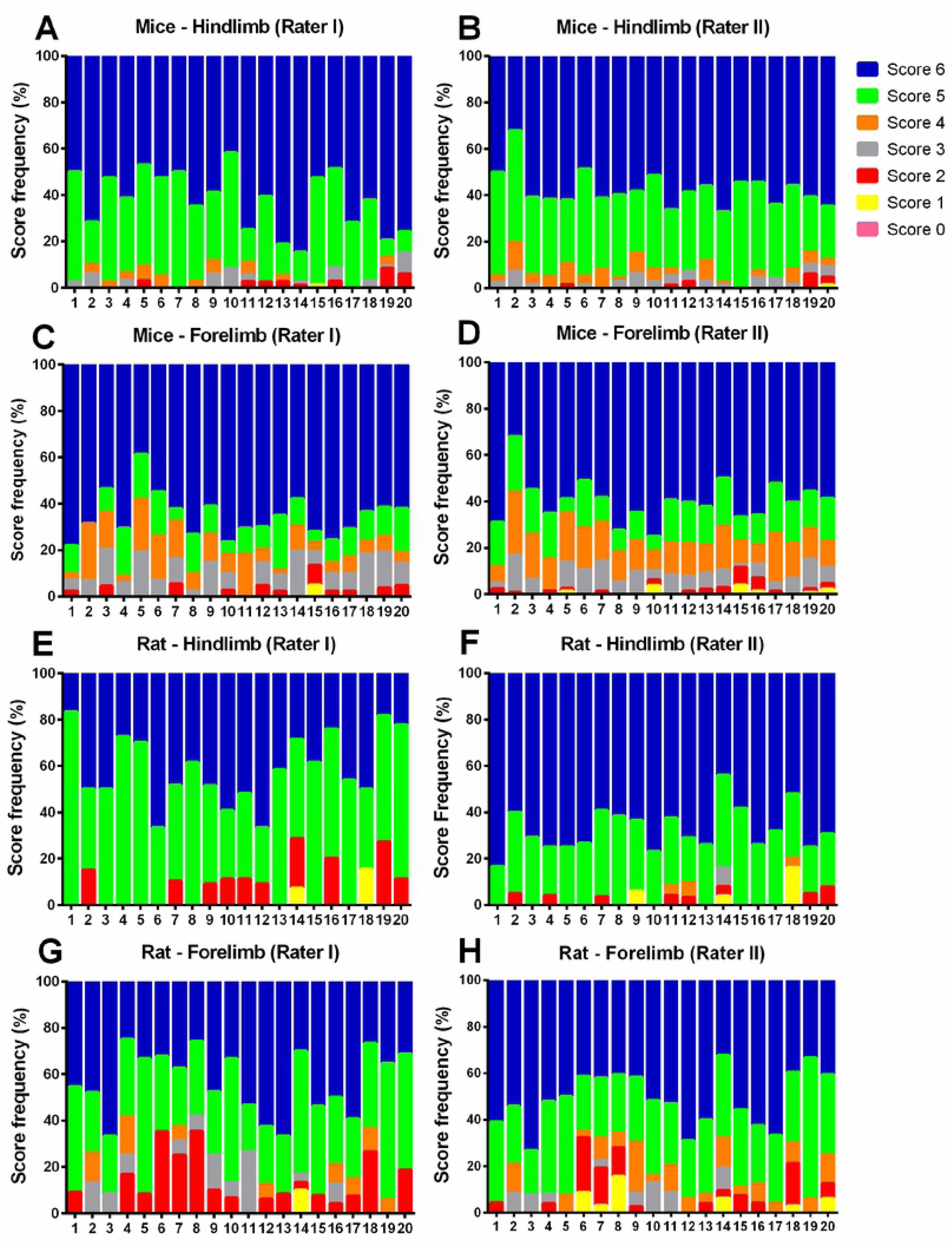
Frequency of scores in the foot fault score (%).

**Figure 5.**
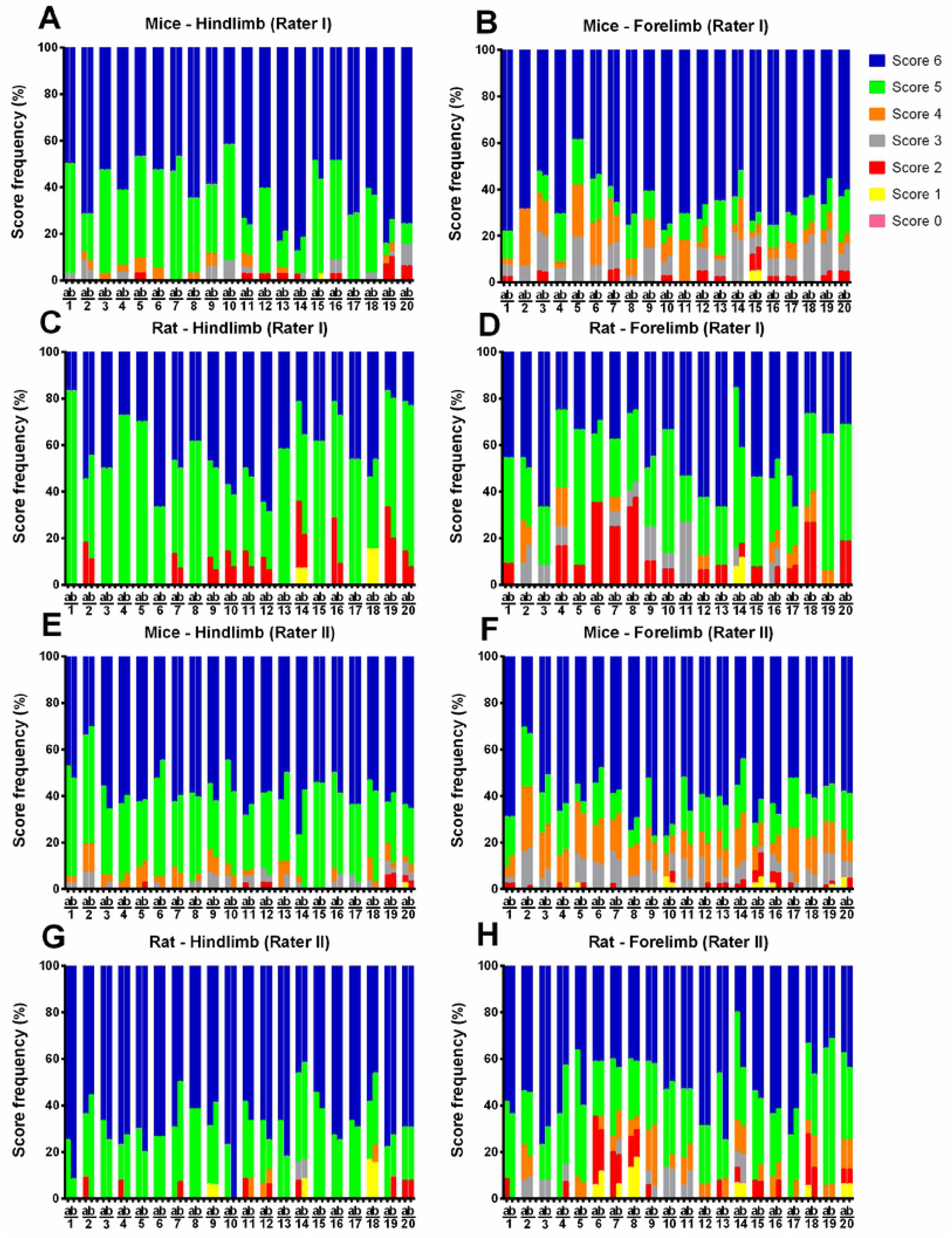
Frequency of scores in the foot fault score (%) in the first (a) and second assessment (b) by Rater I and Rater II.

### 3.2 Inter-rater reliability for mice

The inter-rater reliability score system for mice is shown in Table 3. We observed a strong agreement between the raters in the combined total score (ICC=0.954/P=0.0001), total crossing time (ICC=1/P=0.0001), number of steps (ICC=0.922/P=0.0001) and total time stopped (ICC=0.998/P=0.0001). In addition, the forelimb and hindlimb placement scores showed excellent agreement in the LRWT, with less consistency for forelimb placement (score 3) (ICC=0.466/P=0.090) and hindlimb correction (score 4) (ICC=0.484/p=0.079). Overall, the total scores for the forelimb (ICC=0.925/p=0.0001) and hindlimb (ICC=0.919/p=0.0001) placement between raters I and II showed strong agreement.

**Table 3.** Agreement between raters I and II regarding the outcomes (scores) recorded in the LRWT in mice.

| **Foot fault scoring in mice** |  |  |  |
| --- | --- | --- | --- |
| **Outcome** | **ICC (IC95%)** | **Cronbach's Alpha** | **P Value** |
| Combined Total Score | 0.954(0.883-0.982) | 0.954 | 0.0001* |
| Total Crossing Time | 1(0.999-1) | 1 | 0.0001* |
| Number of Stops | 0.922(0.802-0.969) | 0.922 | 0.0001* |
| Total Time Stopped | 0.998(0.995-0.999) | 0.998 | 0.0001* |
| **Forelimb Placement** |  |  |  |
| **Outcome** |  |  |  |
| Score 0 | 0.889(0.719-0.956) | 0.889 | 0.0001* |
| Score 1 | 0.755(0.381-0.903) | 0.755 | 0.002* |
| Score 2 | 0.699(0.239-0.881) | 0.699 | 0.006* |
| Score 3 | 0.466(0.000-0.789) | 0.466 | 0.090 |
| Score 4 | 0.904(0.757-0.962) | 0.904 | 0.0001* |
| Score 5 | 0.830(0.571-0.933) | 0.830 | 0.0001* |
| Score 6 | 0.712(0.271-0.886) | 0.721 | 0.005* |
| Total Score | 0.925(0.812-0.970) | 0.925 | 0.0001* |
| **Hindlimb Placement** |  |  |  |
| **Outcome** |  |  |  |
| Score 0 | 0.822(0.550-0.929) | 0.822 | 0.0001* |
| Score 1 | 0.889(0.719-0.956) | 0.889 | 0.0001* |
| Score 2 | 0.938(0.844-0.976) | 0.938 | 0.0001* |
| Score 3 | 0.751(0.371-0.901) | 0.751 | 0.002* |
| Score 4 | 0.484(0.000-0.796) | 0.484 | 0.079 |
| Score 5 | 0.622(0.046-0.850) | 0.622 | 0.0001* |
| Score 6 | 0.764(0.405-0.907) | 0.764 | 0.001* |
| Total Score | 0.919(0.795-0.968) | 0.919 | 0.0001* |

\* Statistically significant difference.

### 3.3 Intra-rater reliability for rat

Table 4 shows the intra-rater analyses in rat. We found excellent agreement in the combined total score, total crossing time, number of stops and total time stopped for both raters. Regarding score evaluation, rater I obtained excellent agreement in all the scores for the forelimb (ICC between 0.899 to 0.989 / p=0.0001). Rater II achieved excellent agreement in all scores for forelimb (ICC between 0.787 to 0.920), except for score 6, which was considered satisfactory (ICC=0.652 / p=0.13).

**Table 4.** Intra-rater agreement on outcomes in the analyses of the LRWT in rat.

| **Outcome** | **Rater I** |  |  | **Rater II** |  |  |
| --- | --- | --- | --- | --- | --- | --- |
|  | **ICC (IC95%)** | **Cronbach's Alpha** | **P Value** | **ICC (IC95%)** | **Cronbach's Alpha** | **P Value** |
| Combined Total Score | 0.969(0.922-0.988) | 0.969 | 0.0001* | 0.950(0.875-0.980) | 0.950 | 0.0001* |
| Total Crossing Time | 0.993(0.982-0.997) | 0.993 | 0.0001* | 0.981(0.953-0.993) | 0.981 | 0.0001* |
| Number of Stops | 0.950(0.873-0.980) | 0.950 | 0.0001* | 0.915(0.786-0.966) | 0.915 | 0.0001* |
| Total Time Stopped | 0.806(0.509-0.923) | 0.806 | 0.0001* | 0.939(0.847-0.976) | 0.939 | 0.0001* |
| **Forelimb Placement** |  |  |  |  |  |  |
| Score 0 | 0.899(0.719-0.956) | 0.899 | 0.0001* | 0.919(0.796-0.968) | 0.919 | 0.0001* |
| Score 1 | 0.889(0.719-0.956) | 0.889 | 0.0001* | 0.842(0.600-0.937) | 0.842 | 0.0001* |
| Score 2 | 0.989(0.973-0.996) | 0.989 | 0.0001* | 0.877(0.688-0.951) | 0.877 | 0.0001* |
| Score 3 | 0.978(0.944-0.991) | 0.978 | 0.0001* | 0.920(0.797-0.968) | 0.920 | 0.0001* |
| Score 4 | 0.941(0.851-0.977) | 0.941 | 0.0001* | 0.849(0.618-0.940) | 0.849 | 0.0001* |
| Score 5 | 0.948(0.869-0.979) | 0.948 | 0.0001* | 0.787(0.462-0.916) | 0.787 | 0.001* |
| Score 6 | 0.905(0.761-0.963) | 0.905 | 0.0001* | 0.652(0.121-0.862) | 0.652 | 0.13 |
| Total Score | 0.916(0.787-0.967) | 0.916 | 0.0001* | 0.875(0.685-0.951) | 0.875 | 0.0001* |
| **Hindlimb Placement** |  |  |  |  |  |  |
| Score 0 | 1(1-1) | 1 | 0.0001* | 1(1-1) | 1 | 0.0001* |
| Score 1 | 1(1-1) | 1 | 0.0001* | 0.962(0.904-0.985) | 0.962 | 0.0001* |
| Score 2 | 0.838(0.591-0.936) | 0.838 | 0.0001* | 0.829(0.567-0.932) | 0.829 | 0.0001* |
| Score 3 | 1(1-1) | 1 | 0.0001* | 1(1-1) | 1 | 0.0001* |
| Score 4 | 1(1-1) | 1 | 0.0001* | 0.904(0.758-0.962) | 0.904 | 0.0001* |
| Score 5 | 0.992(0.980-0.997) | 0.992 | 0.0001* | 0.637(0.83-0.856) | 0.637 | 0.16 |
| Score 6 | 0.982(0.954-0.993) | 0.982 | 0.0001* | 0.810(0.519-0.925) | 0.810 | 0.0001* |
| Total Score | 0.988(0.970-0.995) | 0.988 | 0.0001* | 0.970(0.924-0.988) | 0.970 | 0.0001* |

\* Statistically significant difference.

In relation to hindlimb agreement, rater I obtained a similar excellent degree of agreement to that for the forelimb, ranging from ICC 0.838 to 1 / p=0.0001. Whilst rater II achieved a lower agreement than rater I, the ICC was very good, ranging from 0.637 to 1, with only score 5 graded as satisfactory (ICC 0.637). Moreover, both raters obtained excellent intra-rater scores in the outcomes: combined total score, total crossing time, number of steps, total time stopped and total score for forelimb and hindlimb, ranging from ICC 0.806 to 0.993 for rater I and ICC 0.915 to 0.981 for rater II.

### 3.4 Intra-rater reliability for mice

Overall, the intra-rater reliability for mice was excellent for both raters (Table 5). For rater I, in the forelimb foot placement agreement for all the 7 scores were excellent (ICC 0.939 to 1 / p=0.0001). For rater II, the agreement was also excellent, varying between ICC 0.778 and 0.968 for scores 0 to 5. However, score 6 was considered satisfactory (ICC 0.488 / p=0.077). Regarding the hindlimb placement, similar results were found, with the raters only differing in score 6 (rater II obtained a lower ICC: 0.749 / p=0.002) (Table 5).

**Table 5.** Intra-rater agreement on outcomes in the analyses of the LRWT in mice.

| **Outcome** | **Rater I** |  |  | **Rater II** |  |  |
| --- | --- | --- | --- | --- | --- | --- |
|  | **ICC (IC95%)** | **Cronbach's Alpha** | **P Value** | **ICC (IC95%)** | **Cronbach's Alpha** | **P Value** |
| Combined Total Score | 0.971(0.926-0.988) | 0.971 | 0.0001* | 0.963(0.906-0.985) | 0.963 | 0.0001* |
| Total Crossing Time | 1(1-1) | 1 | 0.0001* | 0.999(0.998-1) | 0.999 | 0.0001* |
| Number of Stops | 0.948(0.868-0.979) | 0.948 | 0.0001* | 0.774(0.429-0.911) | 0.774 | 0.001* |
| Total Time Stopped | 0.988(0.969-0.995) | 0.988 | 0.0001* | 0.985(0.963-0.994) | 0.985 | 0.0001* |
| **Forelimb Placement** |  |  |  |  |  |  |
| Score 0 | 1(1-1) | 1 | 0.0001* | 0.919(0.796-0.968) | 0.919 | 0.0001* |
| Score 1 | 1(1-1) | 1 | 0.0001* | 0.778(0.440-0.912) | 0.778 | 0.001* |
| Score 2 | 0.979(0.947-0.992) | 0.979 | 0.0001* | 0.829(0.568-0.932) | 0.829 | 0.0001* |
| Score 3 | 0.979(0.948-0.992) | 0.979 | 0.0001* | 0.899(0.746-0.960) | 0.899 | 0.0001* |
| Score 4 | 0.939(0.846-0.976) | 0.939 | 0.0001* | 0.956(0.888-0.982) | 0.956 | 0.0001* |
| Score 5 | 0.982(0.965-0.995) | 0.982 | 0.0001* | 0.928(0.817-0.971) | 0.928 | 0.0001* |
| Score 6 | 0.950(0.873-0.980) | 0.950 | 0.0001* | 0.488(0.000-0.797) | 0.488 | 0.077 |
| Total Score | 0.978(0.944-0.991) | 0.978 | 0.0001* | 0.934(0.833-0.974) | 0.934 | 0.0001* |
| **Hindlimb Placement** |  |  |  |  |  |  |
| Score 0 | 0.919(0.796-0.968) | 0.919 | 0.0001* | 1(1-1) | 1 | 0.0001* |
| Score 1 | 0.889(0.719-0.956) | 0.889 | 0.0001* | 0.889(0.719-0.956) | 0.889 | 0.0001* |
| Score 2 | 0.978(0.945-0.991) | 0.978 | 0.0001* | 0.963(0.908-0.986) | 0.936 | 0.0001* |
| Score 3 | 0.936(0.839-0.975) | 0.936 | 0.0001* | 0.821(0.548-0.929) | 0.821 | 0.0001* |
| Score 4 | 1(1-1) | 1 | 0.0001* | 0.886(0.713-0.955) | 0.886 | 0.0001* |
| Score 5 | 0.982(0.953-0.993) | 0.982 | 0.0001* | 0.861(0.649-0.945) | 0.861 | 0.0001* |
| Score 6 | 0.958(0.895-0.983) | 0.958 | 0.0001* | 0.749(0.367-0.901) | 0.749 | 0.002* |
| Total Score | 0.950(0.873-0.980) | 0.950 | 0.0001* | 0.951(0.876-0.981) | 0.951 | 0.0001* |

\* Statistically significant difference.

## 4. DISCUSSION

Studying walking adaptability in rodents is of importance to translational neuroscience, since the irregular distribution of the rungs in the walking path requires the animal’s capacity to adjust its stride length, paw placement and control the center of mass. These adaptive motor control strategies are also found and widely studied in humans [28]. Rodents and humans perform some similar movements to protect an injured limb and/or prevent falls [29]. The ladder rung walking test fulfils the fundamental principles of walking adaptability such as pattern of rhythmic reciprocal limb movement; supporting body balance against gravity; and adapting locomotion in response to environmental challenges [1].

Metz and Whishaw created the ladder rung walking test in 2002 to assess forelimb and hindlimb stepping, placing, and coordination in models of cortical and subcortical injury. According to the authors, the test is a sensitive skilled task for assessing slight impairments of walking function and is useful when assessing functional recovery following brain or spinal cord injury and the effectiveness of rehabilitative therapies [14, 30]. Locomotion during the ladder rung walking test is known to depend on ascending and descending neural pathways, since accurately crossing the rungs requires finely adjusted motor control, balance, limb coordination and muscle control [7, 13, 14].

However, to determine the psychometric properties of behavioral tests it is essential to obtain reliable, consistent and scientifically valid findings [31]. Both, intra- and inter-rater agreement are important metrics to ensure reliability and reproducibility [32]. Here, we sought to assess intra- and inter-rater agreement in the foot fault score of the ladder rung walking test using two strains of rodents – Wistar rats and C57BL/6 mice. Two independent researchers (with and without previous experience using the test’s scoring system) analyzed the videos. Our findings suggest the foot fault score system of the ladder rung walking test is a useful, reliable and consistent tool for studying skilled walking performance in rodents. We also found excellent inter and intra-rater reliability for “total crossing time”, “number of stops” and “total time stopped”. The agreement measures provided by this study suggest data obtained by different research groups using the ladder rung walking test should be comparable [33] and encourage the use of the test in further studies.

The ladder rung walking test is an interesting option for researchers investigating neural mechanisms involved in the ability to adapt walking [7, 13, 15, 34]. Since the score reflects the animal’s ability to adapt limb placement and position in a contextual environment [14, 29], the foot fault score system is useful to study walking adaptability in rodents [7, 13]. Whilst traditional biomechanical models of walking analysis require expensive devices, constant animal handling for placing reflective markers and development of signal-processing routines [35, 36], the ladder rung walking test provides walking adaptability assessment using a fast, simple and inexpensive method.

Whereas we observed satisfactory to excellent intraclass correlation indexes in rating individual scores (0 to 6), caution is necessary when using the foot fault score system. Individual scores present subtle differences that may confuse untrained raters. For example, differentiating between scores 3 (replacement) and 4 (correction) requires attention to identify whether the rodent touched the rung before completing paw placement. Moreover, in some situations, the rodent supports a single paw simultaneously on two rungs that are placed too close each other. This may cause confusion in scoring 5 (partial placement) or 6 (correct placement). In addition, rodents sometimes place their paw on the acrylic wall to help walking forward, a behavior that is not considered in the foot fault score system. Furthermore, the subtle differences between score 1 (deep slip) and 2 (slight slip) may cause uncertainty for untrained raters. Finally, the speed of the video recording may also change the perception of the raters during the gait cycle analysis [37]. Thus, the present results suggest experienced and inexperienced raters can get reliable results if appropriate training is provided. We highly recommend the careful study of the article and videos previously published by Metz and Whishaw [14, 15] and supervised practice before using the foot fault scoring system.

Despite being originally designed for rats, the ladder rung walking test can be used in mice with some adjustments to the apparatus, namely a) the diameter of the rungs should be reduced to allow a proper grip and paw placement; and b) the minimal and maximal between-rung interval should be changed, as previously described [13, 24]. Our findings show these adaptations are valid to obtain reliable results in C57BL/6 mice and may be valid for other mice strains.

This study has some limitations. First, only two rodent strains were assessed.

Anyway, the current findings provide evidence of the accuracy and reliability of the foot fault score in both Wistar rats and C57BL/6 mice. Second, we did not compare specific injury models. Despite which, all individual scores (0 to 6) in the foot fault score were found in the studied videos, which minimize this concern.

## 5. CONCLUSION

We conclude the foot fault score of the ladder rung walking test is a reliable and useful tool to study walking adaptability in rodents. Moreover, experienced and inexperienced raters can obtain reliable results if previous supervised training is provided. These findings are of importance for researchers working in the field of translational neuroscience and motor control and impact on the comparability of results obtained worldwide using the foot fault score in the ladder rung walking test.

## ACKNOWLEDGMENTS

We thank Filipe Mello Medeiros and Rodrigo Orso for their valuable contribution in acquiring the video recordings.

